# The Assembly of Purple-red Spinach Provides Insight into the Mechanism of Betalain Biosynthesis

**DOI:** 10.1101/2024.06.06.597681

**Authors:** Zhiyuan Liu, Hongbing She, Meng Meng, Helong Zhang, Zhaosheng Xu, Wei Qian

## Abstract

Purple-red spinach is a unique cultivar with rich in betalains, which has been used for food colorants and is beneficial for human health. In view of the lack of genomic inforamtion of purple-red spinach genome and promote the idetification of betalian gene, we presented the genome of purple-red spinach with Nanopore ultra-long and illumina sequencing platform. The contig N50 size was 2.2Mb and 99.33% contig sequence was anchored to the six chromosomes, totaling 878.83Mb. The purple-red genome constitues 74.15% repeat sequence and harbors 26020 protein-coding genes. We also comprehensively identified candidate genes in betalain biosynthesis pathway in spinach. Moreover, a combined transcriptomic and BSA analysis uncovered a key betalain gene, FUN_002594, which regulated the color and betalain content of leaf. Our data present a valuable resource for facilitating molecular breeding programs of spinach and shed novel light on unique attributes, as well as the modulation of betalain biosynthesis.

## Main text

Spinach (*Spinacia oleracea* L.)is a highly nutritious and economically leafy vegetable crop, which has been widely cultivated throughout the world(Bhattarai and Shi 2021). China was the world largest spinach producing country which yield 27.52 million tonnes spinach, which account for 91% of world production (FAOSTAT, http://faostat3.fao.org). Spinach is excellent rich with vitamins(C, E), minerals(Fe, Ca), flavonoids and low in calories(Howard et al. 2002). The demands for low-calorie diets and benefits from nutritional and health have brought about substantial increasing of spinach over the past few decades.

In spinach, succulent leaves with its stems are edible organ. Therefore, leaf morphological trait, such as leaf color(green with purple-red), leaf surface texture(smooth with savoy) and leaf shape(serrate with entire), are very important agronomical traits of spinach for breeding and commercial values(Cai et al. 2018; Ma et al. 2016). The red color is more appealing for consumer because of it meet up the general interest in the safety, nutritional and aesthetic aspects. Indeed, the purple-red leaf of spinach is attributed to the presence of betalain, a reactive nitrogen-containing pigments (Mou 2019). The betalains are water-soluble and high antioxidant potentials that can scavenge excessive reactive oxygen (Cai et al. 2003).

Betalains has been classified into two types, yellow betaxanthins and red betacyanins. The biosynthetic pathway of betalain has been studied in several species, such as beets(Bean et al. 2018; Sunnadeniya et al. 2016), *Hylocereus undatus*(Chen et al. 2021b) and *Amaranthus tricolor* (Sarker and Oba 2020; Wang et al. 2023). Several key core enzymes were involved in the pathway. In the first step, arogenate dehydrogenase (ADH) catalyze arogenate to form tyrosine(Lopez - Nieves et al. 2018). Then, tyrosine is hydroxylated to L-3,4-dihydroxyphenylalanine (L-DOPA) by unknown enzyme. L-DOPA is converted to betalamic acid by DODA (4,5-DOPA extradiol dioxygenase). Betalamic acid is a key intermediate forming which can spontaneously condense with amino acids or other amine groups to generate yellow fluorescent betaxanthin pigments or spontaneously condense with cyclo-DOPA to generate red betacyanin pigments(Polturak and Aharoni 2019).

In spinach, some effort have been done to identify the gene and marker in the betalain pathway, while no putative betalain biosynthetic genes was dissected(Cai et al. 2018; Ma et al. 2016). Compared to other Caryophyllales species, such as beets and quinoa, the pathway and gene of betalain remains large unclear in spinach. Although several green leaf spinach whole-genome resources are available, the lack of purple-red spinach genome resources great slowed the identification, molecular breeding, and deep utilization of betalain. Here, we generated ∼56 × illumina reads and ∼60 × ONT reads for high-quality genome, and generated a 878 Mb draft assembly. Gene completeness analysis identified purple-red genomes cotaining >99% complete BUSCOs in Embryophyta OrthoDB v10 (n =1614) (Figure 1A).

**Figure 1.**
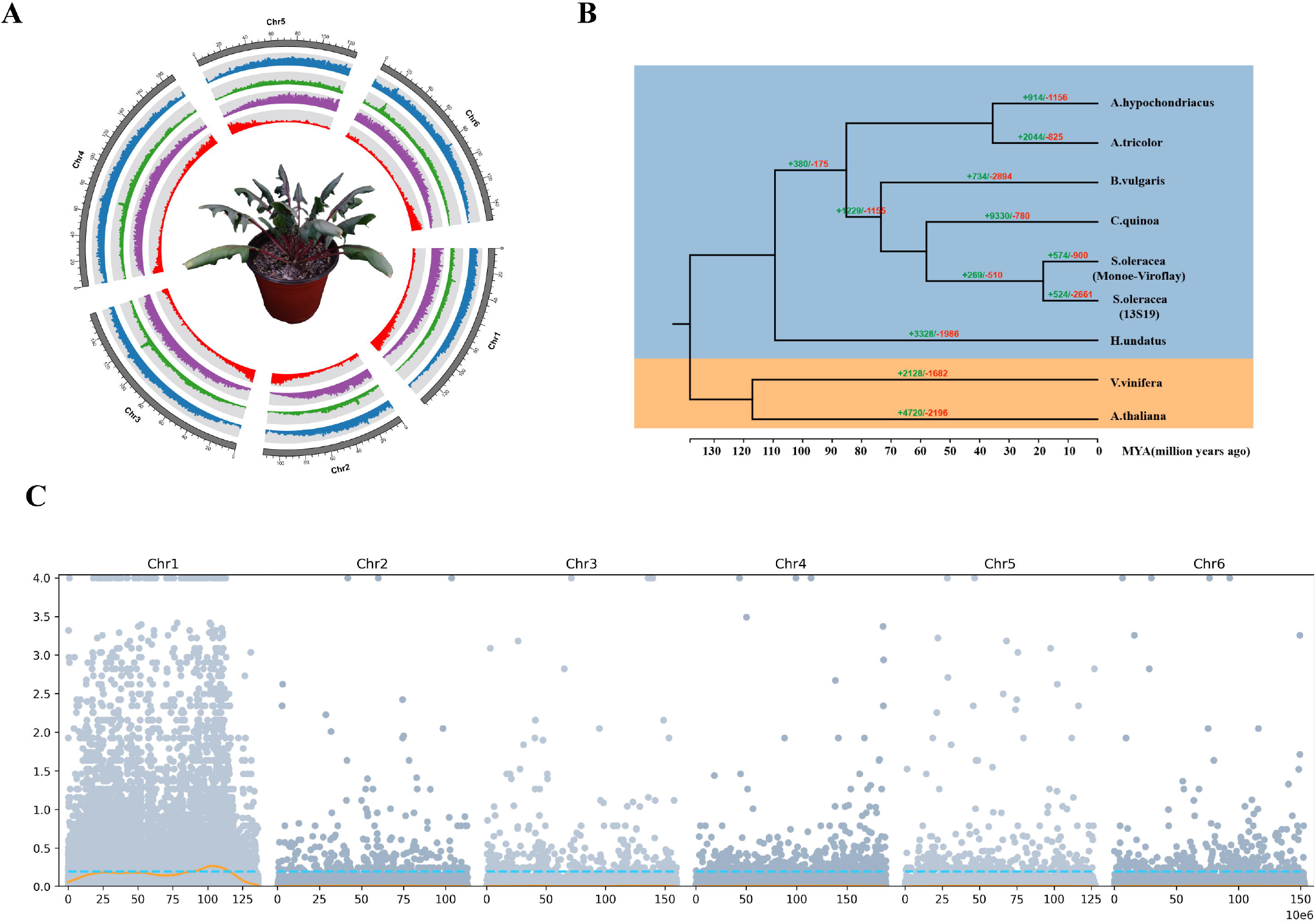
Overview of purple-red spinach genome. (A) The genome feature of purple-red spinach genome assmebly. (B) Species tree and divergene times among nine plants, red and green represent the expanded and contracted gene families, resepectively. (C) Euclidean Distance association analysis.

To explore the genome evolution of purple-red spinach, genes from eight species (*Amaranthus tricolor, Arabidopsis thaliana, Beta vulgaris, Hylocereus undatus, Chenopodium quinoa, Spinacia oleracea*(Monoe-Viroflay), *Vitis vinifera, Amaranthus hypochondriacus*) were clustered into 13945 gene families. Expanded and contracted gene families have been recognized as key drivers that shape the natural variation for adaptation in various species. The CAFÉ program implemented was employed to determine gene families that have undergone considerable expansion or contraction. Comparative genomic analyses were performed among 13S19 and eight representative plant species, 524 and 2661 specific-13S19 gene family expansions and contractions were obtained, respectively (Figure 1B).

The purple-red spinach is renowned for its distinctive purple-red color and unique flavor. To explore the genetic mechanism of purple-red color, we constructed a F_2_ popultaion to analyze the genetic segregation derived from materal line 13S19 and pateral line Sp137. A total of 204 F_2_ individuals were planted, including 158 purle-red and 46 green individuals, respectively. Based on the leaf color of F_1_ (purple-red leaf) and the segregation ratio of 3:1 in F_2_ popultaion, suggesting that the color of leaf was controlling by a single dominant gene. The two bulked pools with purple-red and green leaf were sequenced. The bulk segregant analysis (BSA) identified a major signal on chromosome 1 using ED value(Figure 1C). One region(Chr1: 87.991373-116.03198 Mb) exhibited a higher ED value > 0.268. Subsequently, a large F_2_ population was used to narrow down the region to a 1.6 Mb physical interval. Within this region, we detected a homolog gene of B5GT gene, FUN_002594, named as SpB5GT. We further check the exprssion profile in different color leaf, and found that *SpB5GT* expressed at higher levels in purple-red leaf than in green leaf.

In summary, we present a high-quality purple-red spinach genome of assembly, containg 6 chromosomes. We also identified a key regulating gene of betalain biosynthesis in spinach. These insights offer a deeper understanding of the unique attributes of this specialty cultivar, from its distinctive purple-red color to its special flavor, and the genetic mechanisms underlying these traits.

## Acknowledgments

This work was supported by grants from National Key R&D Program of China (2023YFD1200100), China Agricultural Research System (CARS-23-A-17), Chinese Academy of Agricultural Sciences Innovation Project (CAAS-ASTIP-IVFCAAS).

## Author contributions

Wei Qian conceived and designed the studies. Zhiyuan Liu and Hongbing She performed bioinformatic analyses. Meng Meng prepared samples and conducted RNA-seq analysis. Zhiyuan Liu wrote the draft manuscript. Helong Zhang and Zhaosheng Xu discussed and revised the manuscript. All authors read and approved the final manuscript.

